# A polytypic species revisited: Phylogenetic and morphological variation, taxonomic status, and geographical distribution of *Trachops* (Chiroptera: Phyllostomidae)

**DOI:** 10.1101/2024.01.04.574068

**Authors:** M. Alejandra Camacho, Pablo A. Menéndez-Guerrero, Balázs Horváth, Dániel Cadar, Jérôme Murienne

**Author notes:** **Corresponding author: M. Alejandra Camacho.**, Av. 12 de Octubre, 1076 y Roca. Zip Code: 170525.

## Abstract

The taxonomic status of the neotropical bat genus *Trachops* has been reevaluated through an integrated study that incorporates morphological, morphometric, and molecular data across its extensive geographic range. Our research, which included previously unexamined regions, revealed substantial insights into the diversity within *Trachops*. The results support the elevation of *T. cirrhosus ehrhardti* to species status, due to genetic and morphological differences in southeastern Brazil specimens. Conversely, our comprehensive analysis found insufficient evidence to maintain the subspecific distinction of *T. c. coffini*, which lacks diagnosable morphological characters and is not genetically distinct from *T. c. cirrhosus* across its distribution range. Additionally, our findings challenge the previous belief of a latitudinal differentiation in body size for *Trachops cirrhosus*, as specimens from western South America and northeastern South America exhibit similar sizes to those from Central America. These results underscore the importance of revising the taxonomic framework for this bat genus, contributing to a more precise understanding of its evolutionary relationships, and further enhancing conservation efforts, considering the potential threats to the newly recognized *T. ehrhardti* in the imperiled Atlantic Forest of Brazil.

## Introduction

The Frog-Eating bat, *Trachops cirrhosus* (Spix 1823), is a member of the family Phyllostomidae (Gray 1825), and the only member of its genus (Solari et al. 2019). This species has a wide distribution in the Americas, ranging from southern Mexico to Brazil, and occurring at mid to high elevations at both sides of the Andes Cordillera, across the Amazon, and in the Atlantic Forest (Jones and Carter 1976; Cramer et al. 2001; Williams and Genoways 2008; Solari et al. 2019). The species occurs in humid tropical and subtropical forests, and its distribution encompasses primary, secondary, disturbed and gallery forests, at woodland edge and near cultivated areas (Fenton et al. 1992; Ditchfield 1996; Tirira 2017; Solari et al. 2019).

*Trachops* is monophyletic genus (Ditchfield 1996; Baker et al. 2003; Camacho et al. 2022 and can be easily distinguished from other bats by its finger-like dermal projections on the chin and lips (Figure 1). Historically, *Trachops* taxonomy has undergone numerous changes (Spix 1823; Gray 1825, Gray,1847; Felten 1956a; Schinz 2012) until the late 1950s, when its current taxonomy was stablished with three recognized subspecies: *Trachops cirrhosus cirrhosus* (Spix 1823), occurring from Costa Rica to northeastern Brazil; *Trachops cirrhosus coffini* (Goldman 1925), distributed from Mexico to Nicaragua; and *Trachops cirrhosus ehrhardti* (Felten, 1956b), which is found only in southeastern Brazil (Figure 2) (Jones and Carter 1976; Solari et al. 2019).

**Figure 1.**
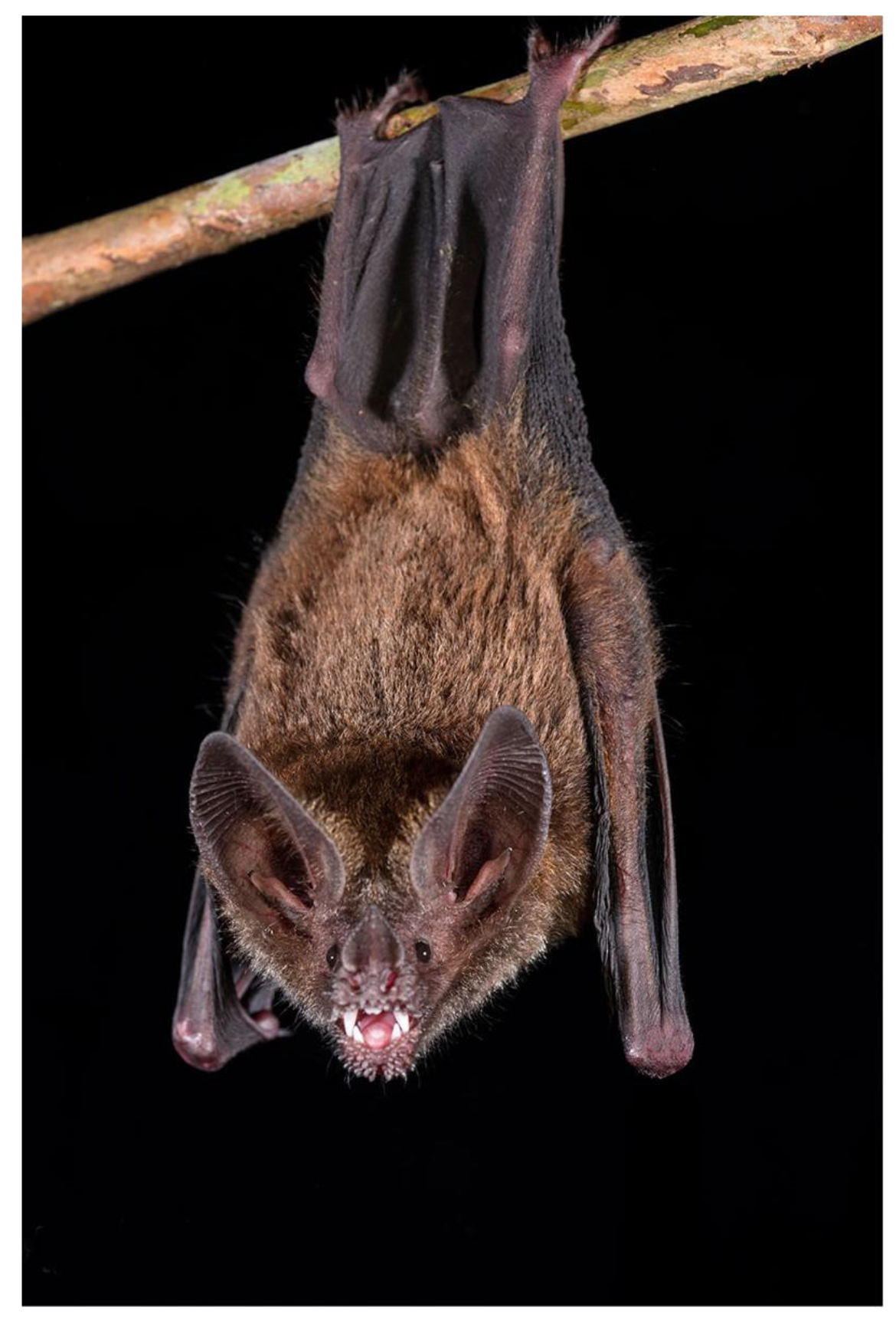
Adult *Trachops cirrhosus* captured in Parque Nacional Yasuní, Orellana, Ecuador. Note the conspicuous finger-like dermal projections on the chin and lips. Photo: Rubén D. Jarrín.

**Figure 2.**
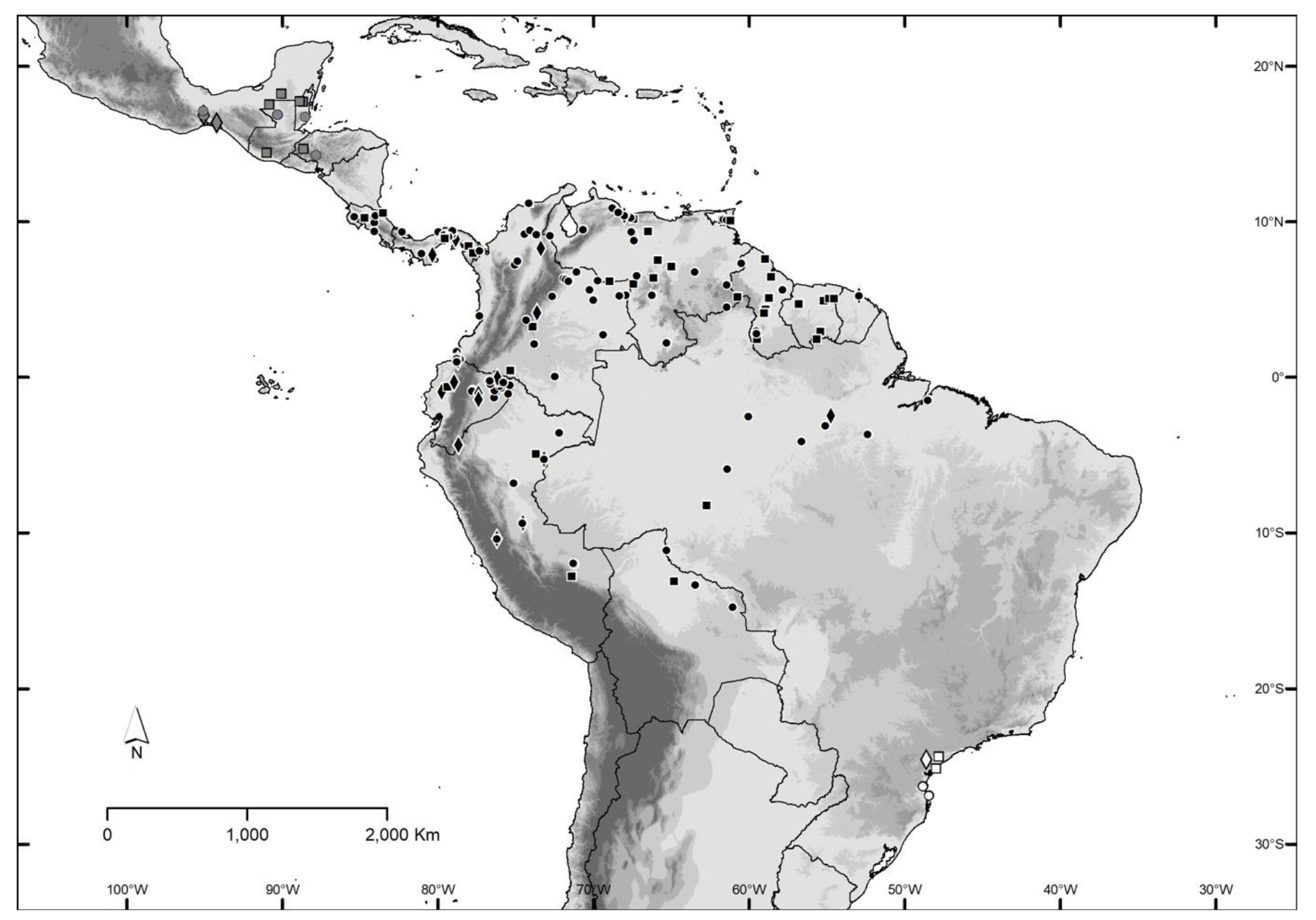
Collecting localities of the *Trachops* specimens analyzed in this study. The localities are shown in the Supplementary Table 1. Black symbols represent *Trachops cirrhosus*, gray symbols represent *Trachops coffini*, and white symbols represent *Trachops ehrhardti.* Circles are specimens that were measured only, squares are sequenced only, and diamonds are both.

The descriptions of the subspecies of *Trachops cirrhosus* are brief and lack comparisons with specimens from Central and South America locations, leading to unclear geographic ranges and diagnostic characters. Moreover, only a few recent studies have addressed the systematics of *Trachops* itself. Ditchfield (1996) used partial sequences of the mitochondrial Cytochrome b gene and identified 5 distinct haplotypes with an average sequence divergence ranging from 5.5% to 11%, demonstrating strong geographic structuring and limited sharing of haplotypes among distant localities. Later, and despite the high genetic divergences yielded by the mitochondrial marker COI, Clare et al. (2007) and Clare (2011) could not find congruence with the topology yielded by the nuclear marker Dby. However, they observed many intraspecific groups occupying sympatric distributions, strongly suggesting ongoing or past speciation events. Most recently, Fonseca (2019) proposed elevating *T. c. ehrhardti* to the species level, based on integrative evidence in morphology, ecology, and genetics, while maintaining *T. c. cirrhosus* and *T. c. coffini* as subspecies of *T. cirrhosus.* But these taxonomic changes, initially proposed by Fonseca (2019), have not yet been formalized in a peer-reviewed publication, and the morphological guidelines for characterization of *Trachops cirrhosus* subspecies remain unclear.

In the present study, we revise the taxonomic composition and phylogenetic structure of the *Trachops* genus based on molecular, morphological, morphometric, and geographic data that includes a significant sample from the West South American region. We aim to address this issue by answering the following questions: (1) Do the current subspecies of *Trachops cirrhosus* have valid phylogenetic and morphological justifications? (2) What are the distinct and diagnosable characters of each subspecies? (3) Can biodiversity of cryptic species be inferred in *Trachops* through molecular and morphological analysis? We seek to clarify, correctly diagnose, and geographically limit *T. cirrhosus* subspecies. We used a phylogenomic dataset based on complete mitochondrial genome which has provided meaningful information in Phylostomids (Camacho et al. 2022). In addition, we performed an extended taxon sampling, ensuring comprehensive spatial coverage. This diverse geographic sampling encompasses a wide range of habitats and ecological zones where *Trachops cirrhosus* subspecies are found.

## Methods

### Specimens, tissue samples, and biorepositories

For the morphometric analyses, we measured 238 specimens of *Trachops cirrhosus*. Only adults were included, following the consideration from Kunz et al. (1996) for sex, age, and reproductive conditions of mammals. We included representatives from all named subspecies and from their complete distribution areas in Central and South America. The samples comprised fluid-preserved specimens, study skins, and skulls deposited in the following institutions: **AMNH**, American Museum of Natural History, New York, NY, USA; **IAvH**, Colección de Mamíferos del Instituto de Investigación de Recursos Biológicos Alexander von Humboldt, Villa de Leyva, Colombia; **MEPN,** Museo de Historia Natural Gustavo Orcés, Escuela Politécnica Nacional, Quito, Ecuador; **QCAZ**, Museo de Zoología, División de Mastozoología, Pontificia Universidad Católica del Ecuador, Quito, Ecuador; **SMF**, Senckenberg Naturmuseum Frankfurt, Frankfurt, Germany; **USNM**, National Museum of Natural History, Smithsonian Institution, Washington, DC, USA; and **UV**, Colección del Mamíferos de la Universidad del Valle, Cali, Colombia. The measurements of the specimens of *Trachops cirrhosus ehrhardti* from the SMF (n = 3) were shared by the curator of this collection. We obtained tissue samples from the following collections: **AMNH**; **FMNH**, Field Museum of Natural History, Chicago, IL, USA; **MSB**, Museum of Southwestern Biology, Albuquerque, NM, USA; **QCAZ**; **ROM**, Royal Ontario Museum, Toronto, Canada; and **SMF** (Supplementary Table 1).

### Genomic DNA isolation, amplification, and sequencing

We gathered a dataset comprising 78 tissue samples, some of them dating back to 1910. The tissue samples encompassed heart, liver, claws, and wing snippets. Mitochondrial DNA sequencing was achieved using a genome skimming procedure following Camacho et al. (2022). Laboratory procedures were carried out at the NGS Core Facility of the Bernhard Nocht Institute for Tropical Medicine in Hamburg. The extraction and amplification of archival DNA were performed in a dedicated clean room facility, which was separate from the area where current samples and post-PCR products were handled. Stringent contamination prevention protocols and negative controls were also implemented.

The DNA extraction process for various sample types (dried skin, ethanol, or formaldehyde-preserved tissues) involved proteinase K digestion at 55°C using 20 µl of proteinase K and 220 µl of ATL lysis buffer (MinElute Reaction Cleanup kit, Qiagen). Prior digestion, the samples were thoroughly washed with nuclease-free water (Qiagen). The incubation time for proteinase K digestion varied depending on the tissue type and sample preservation, ranging from 5 to 24 hours. Following digestion, DNA was extracted and purified using the Qiagen MinElute kit, with each sample eluted to a final volume of 60 µl. DNA concentration was measured using Qubit and Bioanalyzer instruments. For library preparation, the QIAseq FX DNA Library Kit (Qiagen) was used, with double index barcode labelling according to the manufacturer’s instructions. DNA fragmentation was often avoided due to the high degradation of nucleic acid material and low concentration. The HiFi PCR Master Mix from the QIAseq FX kit was utilized to amplify DNA regions with varying GC contents, minimizing sequencing bias caused by PCR, such as nucleotide misincorporations from cytosine deamination. Subsequently, the libraries underwent quality control to determine fragment size using the Agilent 2100 Bioanalyzer, and concentration was assessed using a Qubit 2.0 Fluorometer. After normalization, the samples and negative controls were pooled and subjected to sequencing on the NextSeq 2000 platform (2 × 100 cycles) (Illumina, San Diego, CA, USA).

### Mitogenomes assembly and annotations

Raw reads were first subjected to a qualitative assessment, followed by the removal of adaptor sequences and the filtration of polyclonal and low-quality reads (<55 bases long) using CLC Workbench (Qiagen). Overlapping paired-end (PE) reads were merged to improve quality, while non-overlapping pairs and orphan reads were left unchanged. Deduplication was performed with an assumed 100% identity using BBTools (Bushnell 2014), expanding the length of contigs produced during de novo assembly. Custom assembly was conducted using Megahit (Li et al. 2016) and Spades (Prjibelski et al. 2020) applications. A specialized Chiroptera mitochondrial database and BLASTN were employed to identify potential bat mitochondrial genomes in the resulting contigs. This process was followed by remapping and visual validation of circularization using CLC Genomics Workbench 22. Finally, the GeSeq online tool (Tillich et al. 2017) was used to annotate genomic features. All the assembled mitochondrial genomes were annotated using the MITOS2 metazoan pipeline (Berntet al. 2013; Al Arab et al. 2017), followed by manual adjustment in Geneious v.9.0.5 (https://www.geneious.com). The mitochondrial DNA sequences obtained in this study have been deposited in GenBank (Appendix 1).

### Phylogenetic analysis

Ribosomal RNA (rRNA) and transfer RNA (tRNA) loci were aligned using MUSCLE (Edgar 2004), while protein-coding genes sequences were aligned using TranslatorX (Abascal et al. 2010). As outgroups, we selected sequences from *Phyllostomus hastatus*, *Macrophyllum macrophyllum*, and *Phylloderma stenops*, shown as closely related in previous phylogenetic studies (Botero-Castro et al. 2018; Camacho et al. 2022). We used the optimal partitioning scheme of 38 partitions (2 independent partitions for rRNAs and 36 partitions for tRNAs, one partition for each codon for each gene) and a Generalized Time Reversible (GTR) model of substitution rates along with a gamma (G) distribution and a fraction of invariable (I) sites (GTR+G+I).

We performed a Maximum Likelihood (ML) analysis using RAxML-NG (Kozlov et al. 2019), starting from 10 parsimony trees and 10 random trees. Bootstrap support values were obtained using the classical Felsenstein metric (Felsenstein 1985) and transfer bootstrap expectation (Lemoine et al. 2018). Bayesian inference (BI) analyses were performed using MrBayes v.3.2.7 (Ronquist et al. 2012). We partitioned the sequences in 38 sets corresponding to 2 independent partitions for rRNAs and for tRNAs plus 36 partitions, one partition for each codon for each gene, and used the best analytical scheme as evaluated by the AIC. We ran 8 Markov chain Monte Carlo (MCMC) chains for 10 million generations, with default heating values. The sampling frequency was set every 1000 generations, and the first 25 000 samples were discarded as burn-in. A consensus tree was built under the majority rule consensus of all trees obtained in the 8 runs after the burn-in period. We used the ‘sumt’ command to produce summary statistics for trees sampled during a Bayesian MCMC analysis. Posterior probabilities of nodes were regarded as estimators of confidence. Trees were visualized and edited in FigTree v.1.4.4 (http://tree.bio.ed.ac.uk/software/figtree/). We adopted the nodal support criteria established by Moratelli et al. (2017): in the ML analysis, robust support is considered when bootstrap values exceed or equal to 75%, while support is deemed moderate when values range from 50% to 75%. Values equal to or below 50% are indicative of negligible support in our analyses. We calculated uncorrected pairwise (*p*) distances within and among samples of *Trachops cirrhosus* using MEGA 11 (Stecher et al. 2020; Tamura et al. 2021).

### Morphological and morphometric analyses

A total of 27 measurements were taken. Among these, 9 cranial measurements and 4 external measurements were selected for statistical analyses, as indicated by asterisks (*) below; the remaining measurements were employed for descriptive purposes:

*Calcar length (CL):* From the joint with the ankle to the calcar tip.

*Ear length (E):* Intertragic notch of the ear to the outer tip

*\**Forearm length (FA): Distance from the elbow (tip of the olecranon process) to the wrist (including the carpals). This measurement is taken with partially folded wings.

*Hindfoot length (HF):* Distance from the ankle to the tip of the claw.

*\*Metacarpal III (MET-III):* Distance from the joint of the wrist (carpal bones) with the 3rd metacarpal to the metacarpophalangeal joint of 3rd digit.

*\*Metacarpal IV (MET-IV):* Distance from the joint of the wrist (carpal bones) with the 4th metacarpal to the metacarpophalangeal joint of 4th digit.

*\*Metacarpal V (MET-V):* Distance from the joint of the wrist (carpal bones) with the 5th metacarpal to the metacarpophalangeal joint of 5th digit.

*Tail length (T):* Distance from dorsal flexure at base of the tail to the tip of the last caudal vertebra.

*Tibia length (TiL):* Length from the proximal end of the tibia to the distal base of the calcar.

*Total length (TL):* Head and body length excluding tail.

*Weight (W):* Mass in grams.

*Braincase height (BCH):* Height of the braincase, posterior to the auditory bullae from the basioccipital to the sagittal crest.

*\*Breadth across upper molars (M2-M2):* Greatest width of palate across labial margins of the alveoli of M2s.

*\*Breadth of brain case (BB):* Greatest breadth of the globular part of the braincase, excluding mastoid and paraoccipital processes.

*Condylocanine length (CCL):* Distance from the occipital condyles to the anterior border of the of the upper canines.

*\*Condyloincisive length (CIL):* Distance between a line connecting the posteriormost margins of the occipital condyles and the anteriormost point on the upper incisors.

*Coronoid height (COH):* Perpendicular height from the ventral margin of mandible to the tip of coronoid process.

*Dentary length (DENL):* Distance from midpoint of condyle to the anteriormost point of the dentary.

*\*Greatest length of skull (GLS):* Greatest distance from the occiput to the anteriormost point on the premaxilla (including the incisors).

*Mandibular toothrow length (MANDL):* Distance from the anteriormost surface of the lower canine to the posteriormost surface of m3.

*\*Mastoid (process) breadth (MPW):* Greatest breadth across skull, including mastoid processes. *Maxillary toothrow (MTRL):* Distance from the anteriormost surface of the upper canine to the posteriormost surface of the crown of M3.

*\*Molariform toothrow (MLTRL):* Distance from the anteriormost surface of P3 to the posteriormost surface of the crown of M3.

*\*Palatal width at canines (C-C):* Distance between the outermost extremities of the cinguli of upper canines.

*Palatal length (PL):* Distance from the posterior palatal notch to the anteriormost border of the incisive alveoli.

*\*Post orbital constriction breadth (PB):* Least breadth at the postorbital constriction.

*\*Zygomatic breadth (ZB):* Greatest breadth across the zygomatic arches.

External and osteological characteristics were based on, but not limited to the guidelines proposed by Velazco (2005), Tavares et al. (2014), Molinari et al. (2017), and Garbino et al. (2020). Dental nomenclature follows Miller (1907), Freeman (1998), Garbino and Tavares (2018), and Garbino et al. (2020). Skull, dentition, and external characters were measured with digital calipers (to the nearest 0.01 mm). Total length, tail, hindfoot, ear, and body mass were recorded from skin labels and were only used for descriptive assessments (mean, range, and standard deviation).

We log-transformed the variables and ensuring normality via Kolmogorov-Smirnov tests (Sokal and Rohlf 1995); then we performed two Analyses of Variances (ANOVAs) with the 13 morphological variables to determine sex-based differences within subspecies. With no significant differences found (all *p* < 0.05), we combined data from males and females for further analysis. We used Principal Component Analysis (PCA) as a unified statistical approach to investigate morphological variation in *Trachops cirrhosus*, applying it in two distinct contexts:

1. Morphological Variation Among Subspecies: To assess the morphometric divergence among recognized *Trachops* subspecies, we first applied the PCA focusing on taxonomic identification and current distribution ranges. This involved summarizing key characteristics of the dataset to interpret complex relationships and trends in morphological traits (Sokal and Rohlf 1995). Subsequently, PCA scores were subjected to a multivariate analysis of variance (MANOVA) and Post-hoc multiple comparisons (Holm-Bonferroni correction) to detect significant morphometric differences among subspecies (Rice 1989; Irschick and Shaffer 1997).
2. Geographic Variation Analysis: The second application of PCA was directed towards understanding geographic variation patterns. Here, the same PCA was employed, but with the variable being a more specific geographic distribution area, rather than country boundaries. Geographic areas were established as follows, based on Molinari et al (2023):

1. **CAm - Central America**, from Mexico to Panama, delimited by the Atrato-San Juan Depression to the south, in Colombia.
2. **WSAm - West South America**, defined as the western slopes of the northern Andes, including the Colombian Western Cordillera, the Central Colombian Cordillera, and the Pacific coasts of Colombia, Ecuador, and Peru, to the Central Andes, delimited by the Atrato-San Juan Depression to the north and the Southern Andes to the south.
3. **ESAm - East South America**, demarcated as the upper forested slopes on the eastern side of the Andes and the low Amazon of Colombia, Ecuador, Peru, and Bolivia; the entire Amazon drainage of Brazil, including northeastern Bolivia, and the Orinoco drainages of Colombia and Venezuela.
4. **NEC/NWV,** NE Colombia and NW Venezuela, comprising the Orinoco llanos and Andean piedmont plus the Venezuelan Coast range, including the Guajira peninsula and the Maracaibo depression (Ferrer-Pérez et al. 2009).
5. **GS - the Guyana Shield**, including the Venezuelan states of Bolivar and Amazonas, and a portion of Delta Amacuro, the entire territories of Guyana, Surinam, and French Guyana, and parts of northern Brazil.
6. **AF - Atlantic Forest**, confined to the southernmost eastern region of Brazil, including the Mata Atlántica ecosystem.

## Results

### Specimens, tissue samples, and biorepositories

Altogether, we measured 238 specimens of *Trachops cirrhosus* from all named subspecies (i.e., 202 *T. c. cirrhosus*, 31 T*. c. coffini*, and 5 *T. c. ehrhardti*), sourced from 7 scientific collections (Supplementary Table 1). Additionally, out of the 78 tissue samples, we successfully sequenced a total of 54 complete mitochondrial genomes (i.e., 47 *T. c. cirrhosus*; 5 T*. c. coffini*; 2 *T. c. ehrhardti*; Supplementary Table 1, Appendix 1). The remaining 24 individuals that could not be sequenced corresponded, mostly, to samples of ancient tissues preserved in formalin. 22 specimens were both measured and sequenced. To analyze geographic data, we used 173 unique collection locations, duly documented (Figure 2).

### Phylogenetic analysis

Monophyly of *Trachops* was recovered with strong support from both the ML analysis (bootstrap support [BS] = 100%), and from the Bayesian analysis (Bayesian posterior probability [BPP] = 1). There is broad agreement between analysis methods (ML and BI) for this dataset. We found two well-supported and genetically distant clades: one containing the subspecies *T. c. coffini,* nested within *T. c. cirrhosus* (Clade 1; Figure 3) while the other contained the subspecies *T. c. ehrhardti* (Clade 2, Figure 3).

**Figure 3.**
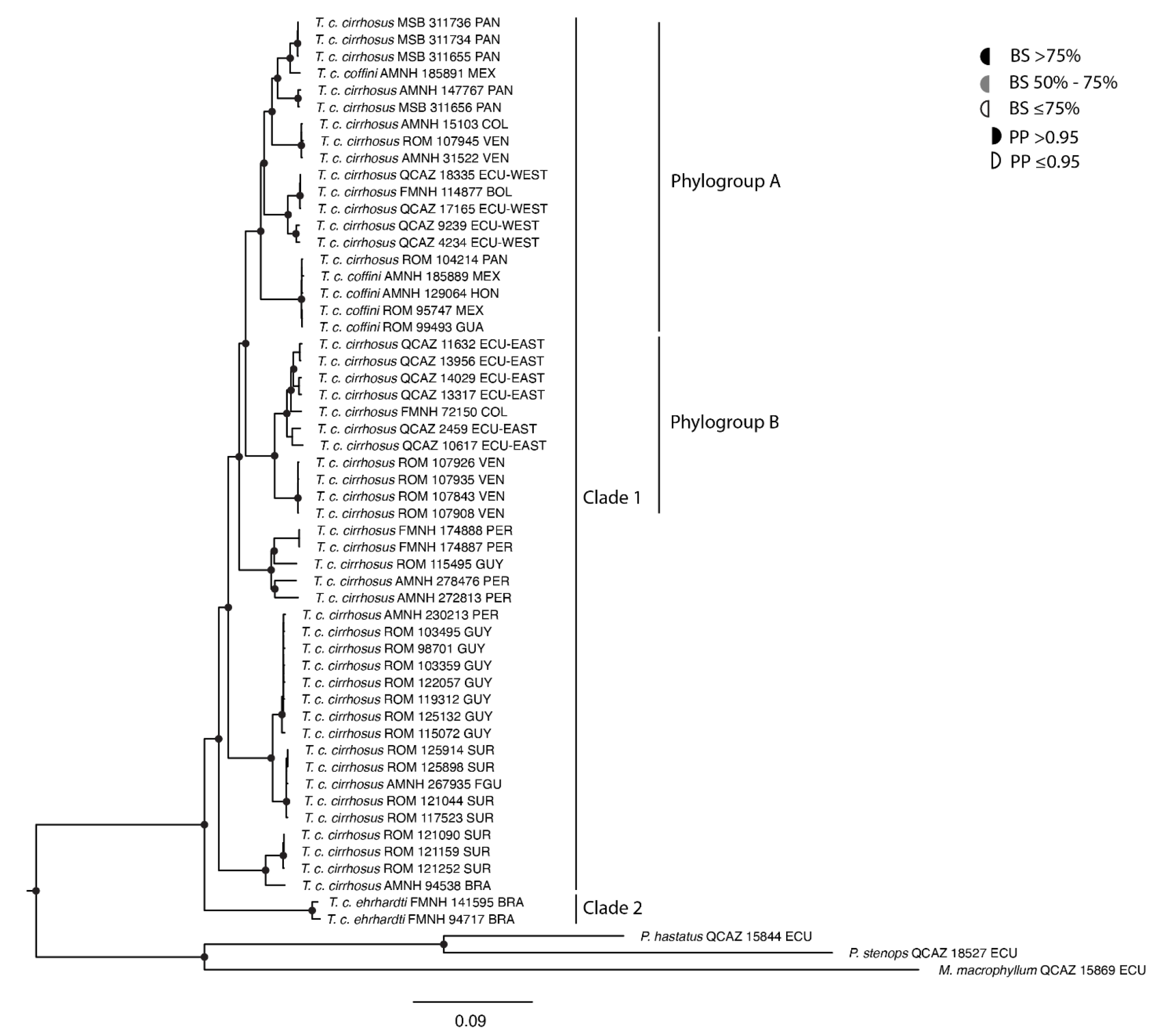
Maximum Likelihood phylogeny of *Trachops* based on complete mitochondrial genomes (nucleotide sequences). The tree was reconstructed in RAxML under the GTR+G+I model using 57 complete mitogenomes including outgroups. Shaded semicircles on the nodes indicate ML bootstrap support (as a percentage) and Bayesian posterior probabilities from a congruent Bayesian Inference Analysis (see insert key).

In Clade 1 we recovered 2 clear phylogroups: Phylogroup A comprising samples from Mexico, Guatemala, Honduras, Panama, western and northeastern Colombia, and northwestern Venezuela, and western Ecuador. According to the geographical groups analyzed in the morphometric analyzes (see Methodology and following section of Results), all these sequences correspond to the geographic groups CAm (Central America), WSAm (West South America) and NEC/NWV (NE Colombia and NW Venezuela), except for one sequence from Bolivia (FMNH 114877), which in the ML topology appears nested with individuals from western Ecuador. Within this phylogroup, there is a sequence from the province of El Petén, Guatemala, which is located 93 km away from the type locality of *T. c. coffini* (Guyo, Petén, Guatemala). The subspecies do not align with geographically distinct monophyletic groups, and furthermore, they do not match the geographical distributions historically associated with *T. c. cirrhosus* and *T. c. coffini* subspecies. Mean pairwise uncorrected sequence distances between what has been named as *T. c. cirrhosus* and *T. c. coffini* is 5.6% (Table 1), which is not different from the computed intraspecific divergence values of 5.5% in *T. cirrhosus* (Table 2).

**Table 1.** Estimates of evolutionary divergence across sequence pairs between groups are presented. The number of base differences per site, calculated by averaging over all sequence pairs between groups, is depicted below the diagonal and scaled as percentages.

|  | *T. c. cirrhosus* | *T. c. coffini* | *T. c. ehrhardti* | Outgroup |
| --- | --- | --- | --- | --- |
| *T. c. cirrhosus* |  | 0.0013 | 0.0018 | 0.0023 |
| *T. c. coffini* | 5.6% |  | 0.0022 | 0.0025 |
| *T. c. ehrhardti* | 8.2% | 8.6% |  | 0.0025 |
| Outgroup | 17.7% | 17.8% | 17.8% |  |

**Table 2.** Estimates of evolutionary divergence across sequence pairs within groups Standard error estimate(s) are shown in the second column and were obtained by a bootstrap procedure (1000 replicates).

|  | Distance % | S.E. (%) |
| --- | --- | --- |
| *T. c. cirrhosus* | 5.5 | 0.0 |
| *T. c. coffini* | 1.9 | 0.0 |
| *T. c. ehrhardti* | 0.0 | 0.0 |
| Outgroup | 0.1 | 0.0 |

We have identified a sister clade to Phylogroup A that we named as B (Figure 3), comprising individuals from Eastern Ecuador and Eastern Venezuela (Orinoco Basin), corresponding to the geographical group ESAm (East South America), as identified in our morphometric analyses. Besides these two strongly supported phylogroups, no further distinct phylogenetic relationships were found within Clade 1, although there is a noticeable clustering of samples from the northeastern region of South America, corresponding to the Guyana Shield.

According to the concordant phylogenies of ML and BI analyses, Clade 2 forms a monophyletic clade containing specimens from the southeastern region of São Paulo, in Brazil, which is geographically close to the state of Santa Catarina (type locality of *T. c. ehrhardti*). According to the geographical groups analyzed in the morphometric analyses (see Methodology and the following Results section), these sequences correspond to the geographic group AF (Atlantic Forest; Figure 3).

### Morphologic and morphometric variation

Our examination of the *Trachops* subspecies highlights the morphological congruence, particularly in qualitative traits, between *T. c. cirrhosus* and *T. c. coffini*, contrasting with the distinct morphology of *T. c. ehrhardti*. Our scrutiny failed to discern marked morphological divergences between *T. c. cirrhosus* and *T. c. coffini*, challenging their current classification as distinct entities. The attributes previously considered distinctive, such as skull shape, rostrum morphology, dimensions of premolars and molars, and mandible features, displayed uniformity across these two subspecies in our geographical sampling (Figure 4). Table 3, illustrating variable ranges among the *Trachops* subspecies, suggests a considerable overlap, rendering these variables ineffective as discrete discriminators among the 3 subspecies.

**Figure 4.**
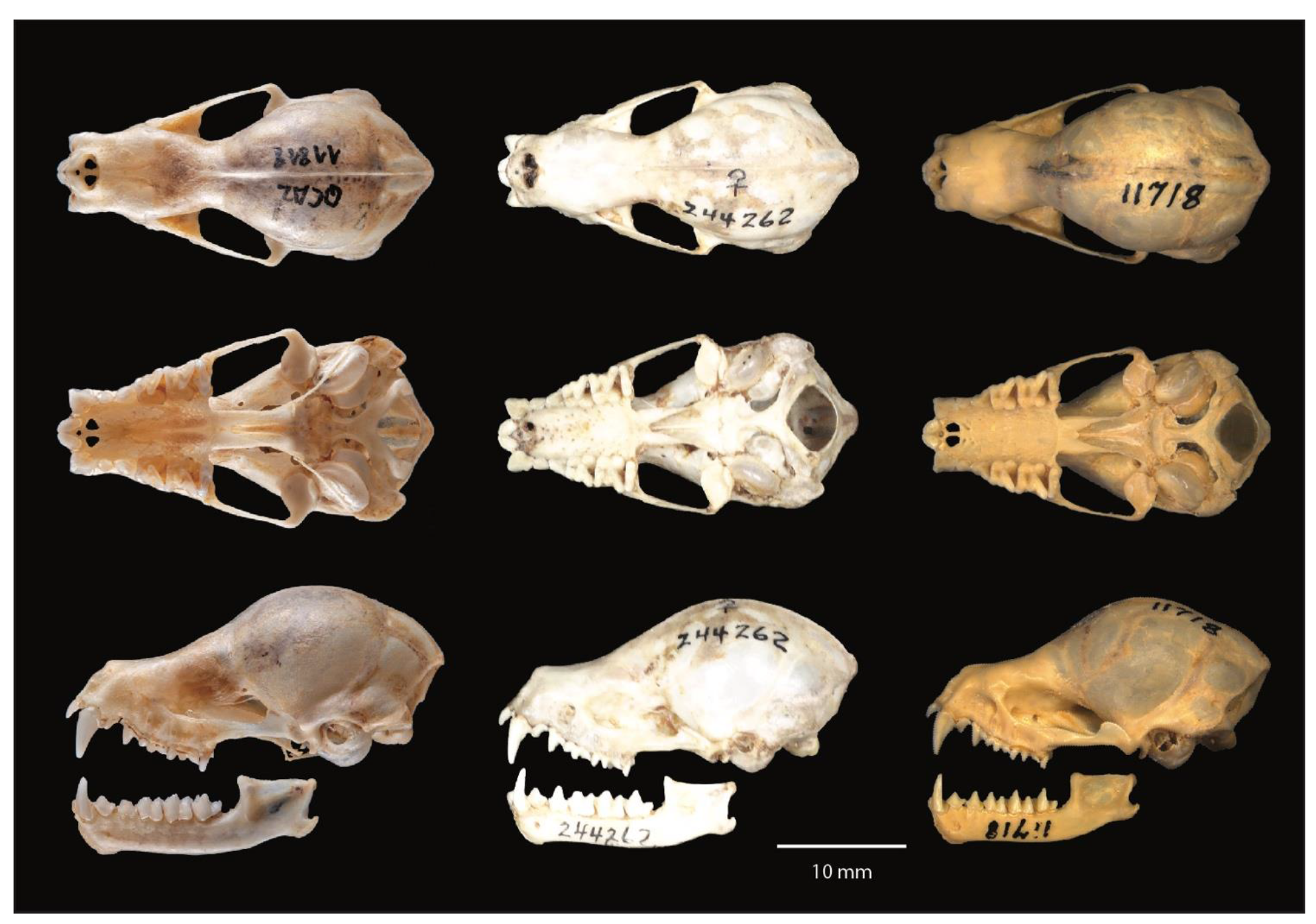
Dorsal, ventral, and lateral views of the skulls and lateral view of the mandibles of *Trachops cirrhosus cirrhosus* (left); *T. c. coffini* (middle), and *T. c. ehrhardti* (right) Photos: Rubén D. Jarrín (QCAZ 11818); M. Alejandra Camacho (NMNH 244262); Anika Vogel (SMF 11718).

**Table 3.**
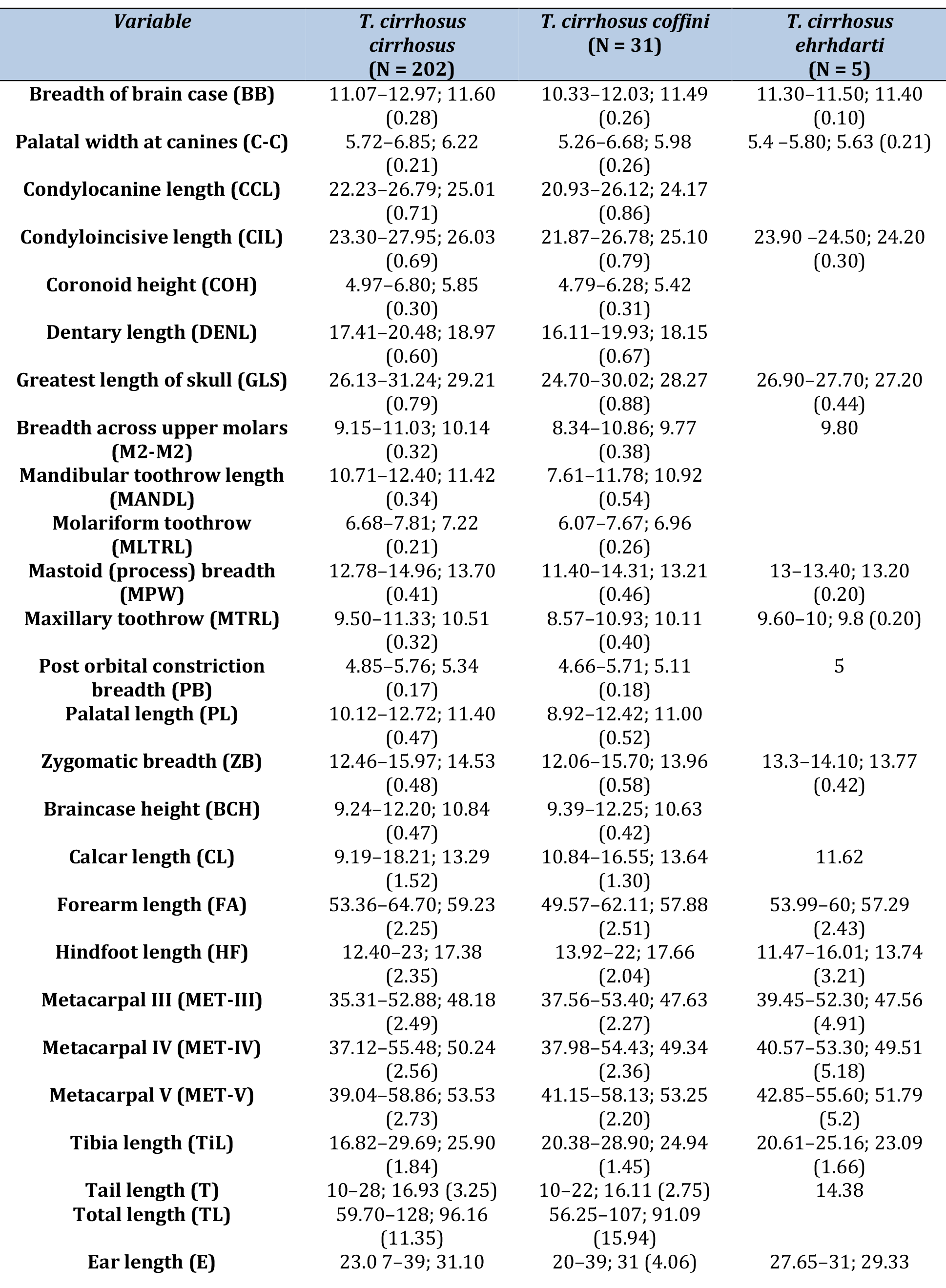
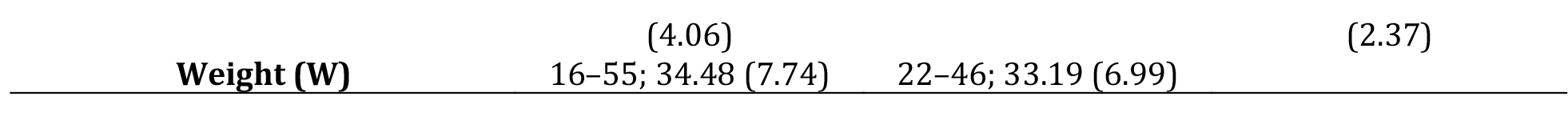
Measurements (mm) and body masses (g) of *Trachops* c. *cirrhosus*, *T. c. coffini* and *T. c. ehrhardti*. Descriptive measures: minimum–maximum; arithmetic mean (standard deviation).

In the Principal Component Analysis (PCA), two principal components cumulatively accounted for 73% of the total variance in our log-transformed dataset (Table 4). The PCA plot (Figure 5) reveals substantial overlap in the 2D morpho-space among the subspecies, attributed to their size and shape congruities. Notably, *T. c. cirrhosus* is positioned towards the right on PC1, indicating its relatively larger size compared to *T. c. coffini*, which is leaned towards the left. However, this positioning does not demarcate distinct groupings. PC1 predominantly shows size variations, influenced by skull length metrics such as Condyle-Incisive Length (CIL), Greatest Length of Skull (GLS), Maxillary Palatal Width (MPW), and Maxillary Toothrow Length (MTRL), each with factor loadings exceeding 0.8. Conversely, PC2 appears to reflect shape variations, primarily influenced by Palatal Breadth (PB).

**Figure 5.**
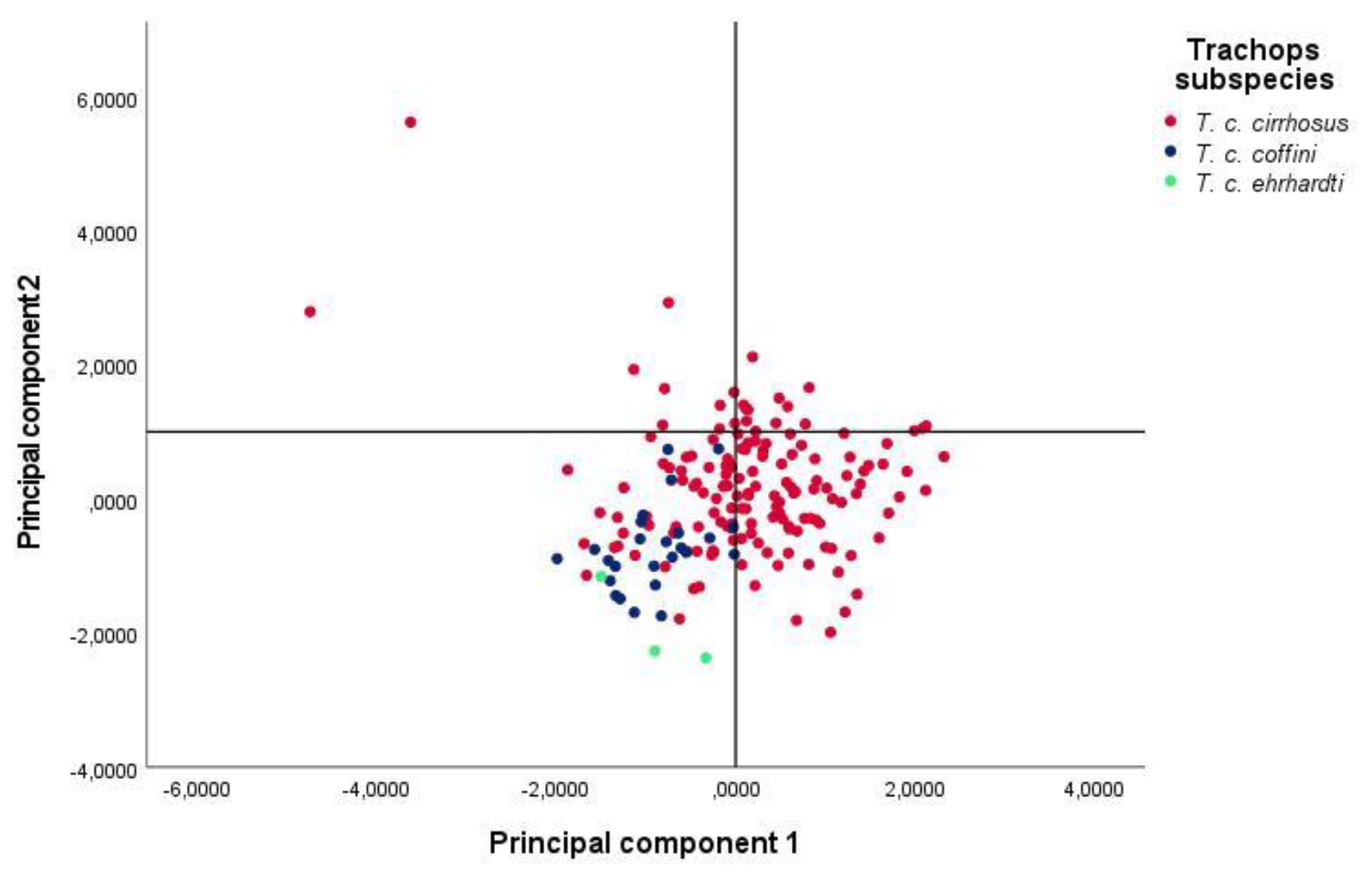
Principal components (PC’s) from a PCA based on 13 measurements from 238 individuals. Samples: *Trachops cirrhosus cirrhosus* (red circles), *T. c. coffini* (blue circles), and *T. c. ehrhardti* (green circles).

**Table 4.**
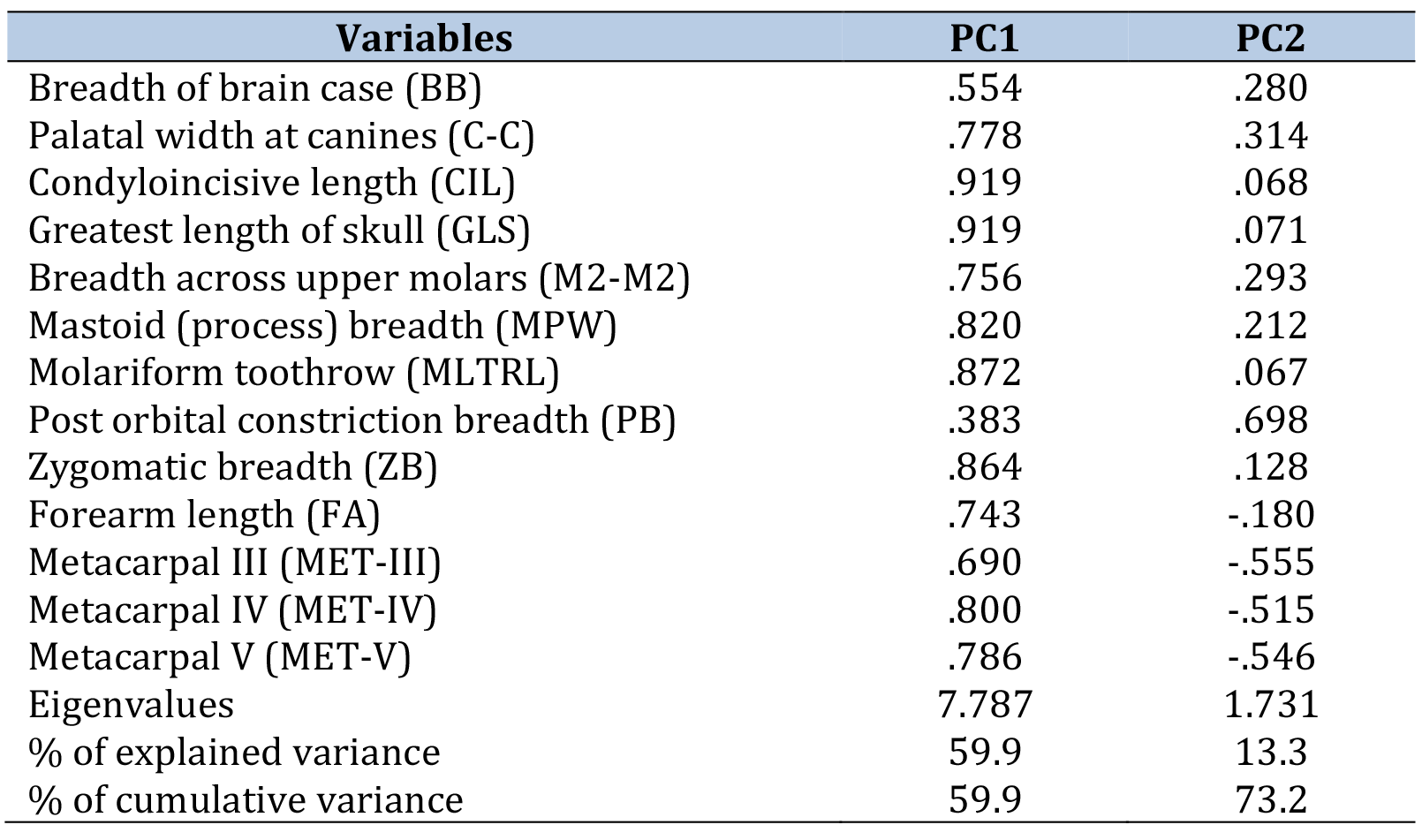
Loadings, eigenvalues and percentage of variance for 2 principal components from a PCA of the 13 linear measurements for adult specimens of *Trachops*.

Despite the apparent overlap in morpho-space depicted in Figure 5, our multivariate analysis of variance (MANOVA), employing Pillai’s Trace test, identified statistically significant differences among the subspecies on both PC1 and PC2 (F = 16.373, *p* < 0.01). Further post-hoc multiple comparison analyses delineated a statistical distinction between *T. c. cirrhosus* and *T. c. coffini* in terms of size on PC1 (*p* < 0.01), but not between *T. c. cirrhosus* and *T. c. ehrhardti* (p > 0.05). Moreover, no statistical differences were noted between *T. c. coffini* and *T. c. ehrhardti* (p > 0.05). These statistical disparities might be influenced by the limited sample size for *T. c. ehrhardti*, which does not sufficiently echo in morphological differentiation. The observed statistical differences, particularly between *T. c. cirrhosus* and *T. c. coffini*, are evident in the context of size, as indicated by PC1. However, the overarching morphological congruence, especially considering the overlap in the PCA plot, suggests that these subspecies, despite some statistical variance, share a degree of morphological commonality.

### Geographic variation

There is a noteworthy overlap in the 2D morpho-space between the geographic-based groups (Figure 6). The multivariate analysis of variance suggested there are differences between the geographic groups on PC1 and PC2, as suggested by Pillai’s Trace test (F = 17.629, *P <* 0.01). Post-hoc multiple comparison analyses unveiled variation in size (PC1) among the samples. Specifically, there was a size difference between the samples from Central America (CAm) compared to those from East South America (ESAm), the Guyana Shield (GS), and North South America (NEC/NWV), but not when compared to the West South American (WSAm) samples. The plot illustrates that both CAm and WSAm samples cluster to the left along PC1, denoting their smaller size, but they do not end up forming a separate cluster. Conversely, the Atlantic Forest (AF) specimens were notably smaller in size compared to Guyana Shield (GS) samples, though not significantly different from the rest of the regions. In fact, the GS sample exhibited the largest size relative to all other groupings. Notably, NEC/NWV specimens did not display a clear grouping pattern. Regarding the PC2, AF specimens exhibited notable differences, particularly in variables such as palate width, in comparison to ESAm specimens. Additionally, a highly significant difference was observed between CAm and ESAm specimens.

**Figure 6.**
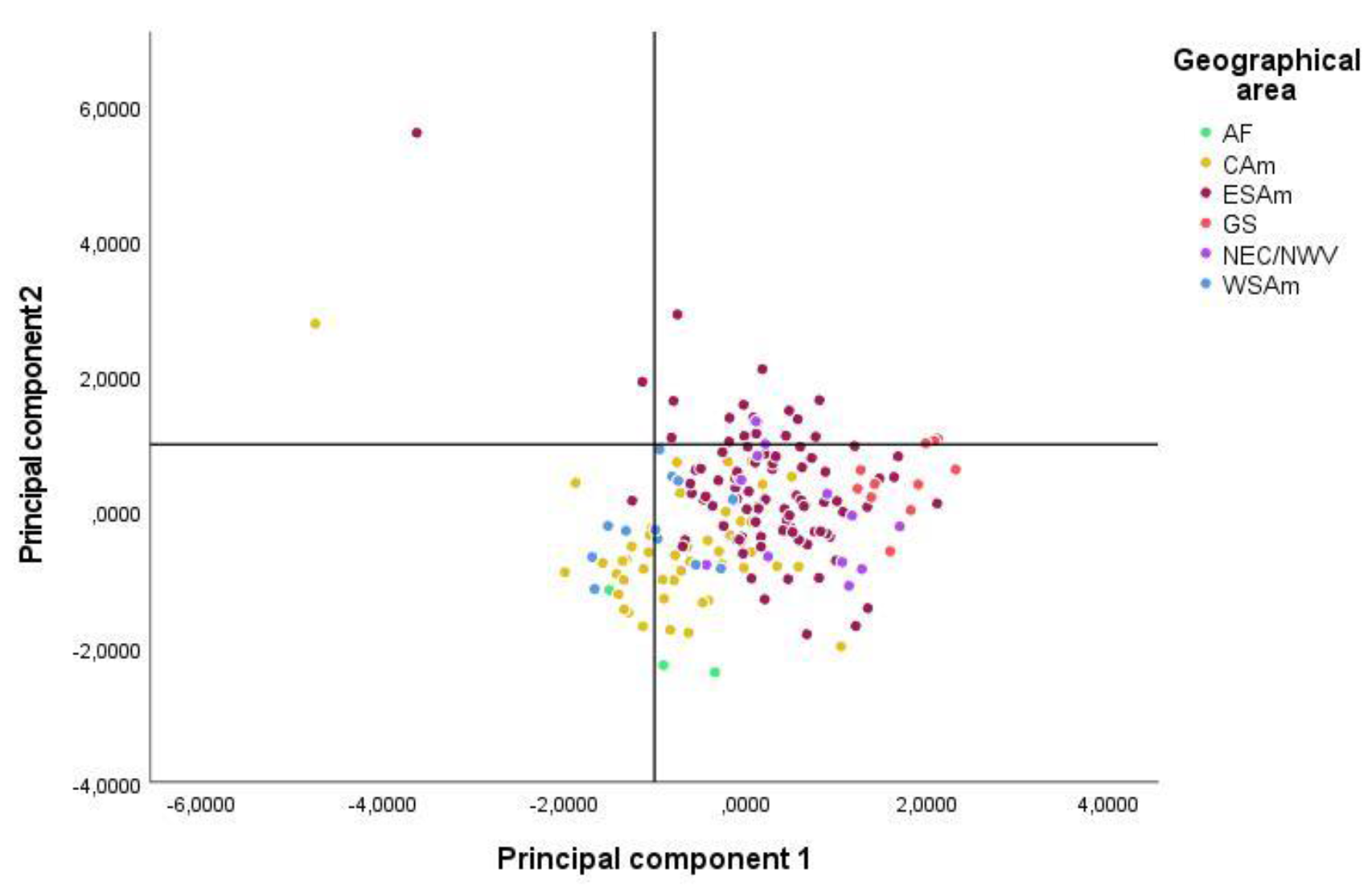
Principal components analysis of 6 geographic groups and 238 specimens of *Trachops* of the 13 cranial measurements considered in the morphometric analysis. Meaning of Geographic abbreviation areas: **CAm** - Central America; **WSAm** - West South America; **ESAm** - East South America; **NEC/NWV**, NE Colombia and NW Venezuela; **GS** - the Guiana Shield; and **AF** - Atlantic Forest.

Both qualitative and quantitative assessments reveal a noticeable degree of morphological and genetic similarity between the traditionally recognized subspecies *T. c. cirrhosus* and *T. c. coffini.* Moreover, our geographical analysis reveals that the distribution of morphological variation among different regions does not align with the historical classification of subspecies. These findings strongly support their consideration as part of the same taxonomic group. Additionally, our investigation provides definitive evidence for elevating *T. c. ehrhardti* to the status of a distinct species, due to significant genetic and morphological disparities observed in specimens from the southeastern region of Brazil.

## Discussion

Our investigation of *Trachops*, which integrates data from morphology, morphometrics, and molecular analysis encompassing almost the totality of its distribution range, has provided clarification into the taxonomic status of the species. Our main findings include the recognition of specific status of *T. ehrhardti,* and *T. c. coffini* as a junior synonym of *T. cirrhosus*. Incorporating *T. cirrhosus* specimens from Colombia, Ecuador, and Peru was crucial to our classification, as it represented the first taxonomic exploration of the species in the West South American region. We not only expanded the geographical scope of the analysis, but also shed light on previously unexplored genetic and morphological variations within *T. cirrhosus* populations.

### Trachops in phylogenetic context

The mitochondrial DNA (mtDNA) diversity in *Trachops cirrhosus* is significant, as evidenced by three previous studies. Using 10 partial sequences of the mitochondrial gene Cytb from localities such as Guatemala, Panama, French Guyana, and Brazil, Ditchfield (1996) identified 5 clades.

These findings suggested that current subspecies classifications may underestimate haplotypic diversity, indicating that *Trachops* should be considered a complex. In this topology, *T. c. coffini* is not monophyletic as opposed to *T. c. cirrhosus,* and the author attributed this haplotypic diversity to reduced mobility or increased philopatry in *Trachops* across its geographic range, though these hypotheses remain unconfirmed (Ditchfield 1996).

Later, Claire (2011) investigated pattern congruence between two independently evolving genetic regions (mitochondrial genome and Y chromosome), finding significant mitochondrial haplotypic divergence, but not in the Dby 7th intron region. Despite using more samples than Ditchfield (1996), the author did not include specimens from West South America. In this topology, *T. c. coffini* was once again not a monophyletic clade distinct from *T. c. cirrhosus*, lacking geographical congruence to support its separation as a subspecies.

Finally, Fonseca (2019) proposed a more integrative approach with a greater number of sequences using a dataset composed of 2341 bp of Cyt-b, COI, D-loop and STAT5A from 129 tissues. Samples from western South America, particularly Colombia, Ecuador, and Peru, were not included in this study. In consistency with Clare et al. (2011), the ML phylogeny did not render *T*. *c. coffini* as an independent clade of *T*. *c. cirrhosus*, with little genetic differentiation (Fonseca 2019). The author proposed that *Trachops* should be divided into two lineages recognized at the species level: *Trachops ehrhardti*, monotypic, and *T. cirrhosus*, with two subspecies (T. *c. cirrhosus* and T. *c. coffini*). However, the phylogenetic justification for maintaining *coffini* as a subspecies and its distribution was unclear.

In this study, we included an extensive taxon sampling with new localities from Colombia, Ecuador, and Peru, as well as information from complete mitochondrial genomes. Our findings revealed two reciprocally monophyletic clades, each exhibiting significant genetic divergence. The first clade, *T. cirrhosus*, notably includes sequences from northern Central America (*T. c. coffini*). This challenges the classical notion of *T. c. coffini* as a distinct subspecies, suggesting instead a more cohesive genetic identity within the *T. cirrhosus* clade. The second clade, *T. ehrhardti*, stands as a monophyletic group, sister to *T. cirrhosus*. These results prompt a reconsideration of *Trachops’* taxonomic structure, particularly the delineation of *T. ehrhardti* and *T. cirrhosus*, and the subspecies status of *T. c. coffini* within this framework.

### Trachops in the morphometric context

When *T. coffini* was described as a separate species by Goldman (1925), he reported larger measurements for *T. cirrhosus*. The author revised 18 specimens from the type locality of *T. coffini* with others from Venezuela, Colombia, and Panama, assuming that they represented typical *T. cirrhosus*, without specifying quantity or localities. However, his analysis, based on a limited geographical range and a small sample size, raised questions about the robustness of these early findings. Felten (1956a) and Felten (1956b) revisited this classification, incorporating a broader geographic range, including Central America. His work, which suggested classifying the Central American forms as *T. c. coffini* subspecies, highlighted the potential intraspecific variation within the species. However, Felten’s conclusions, particularly regarding the dental morphology and size differences, were largely inferential, lacking quantitative rigor. The author proposed maintaining the two subspecies and added *T. c. ehrhardti* as a subspecies, based on 3 specimens from southeastern Brazil, with no apparent difference other than the smaller size, in relation to samples from Colombia, recorded by Hershkovitz (1949), but similar in size to the samples from El Salvador (Felten 1956a).

No other significant morphological study of *Trachops* was published until six decades later, when Fonseca (2019) revised specimens mostly from localities in Brazil. The quantitative analysis revealed a substantial overlap in the morphometric space across the three subspecies studied, notably between *T. c. cirrhosus* and *T. c. coffini*, challenging the notion of distinct morphometric differentiation between these subspecies. This finding was an indicative of a morphometric continuum rather than discrete categories. Qualitatively, she observed that *T. c. coffini*, exhibited a pronounced angle between the rostrum and the braincase in lateral view, a characteristic less evident in *T. c. cirrhosus*. In dental morphology, Fonseca (2019) noted that the first lower premolar (p1) in *T. c. coffini* possesses a unique shape, distinct to this subspecies, although the specifics of this form were not delineated. Additionally, she reported that the m1 tooth in *T. c. coffini* demonstrates a more developed paraconid, compared to *T. c. cirrhosus*. The potential displacement of p3 to the labial surface in *T. c. coffini*, as opposed to the lingual surface in *T. c. cirrhosus*, suggests further morphological variability. Notably, Fonseca (2019) noted that the specimens from Costa Rica displayed morphological traits and cranial sizes more akin to T*. c. coffini*, diverging from the expected distribution patterns. Furthermore, the individuals from Panama exhibited traits from both *T. c. coffini* and *T. c. cirrhosus* in equal frequencies. This observation led Fonseca to propose a clinal variation in size across the species’ range, with smaller individuals observed in Central America and larger ones in South America, positioning Panama as a zone of intergradation.

In our study, encompassing specimens from the western region of the Andes and northern South America, we did not observe the diagnostic characters reported by Fonseca (2019). Specifically, our samples from Central America did not exhibit the morphological differences between the northern and southern regions as previously proposed. This discrepancy, particularly regarding the distinct cranial angles in *T. c. coffini* and the unique shape of p1, suggests a less complex morphological variability within the *Trachops* genus. Even though the Principal Component Analysis (PCA) revealed a size variation on PC1, predominantly influenced by skull length metrics, the PCA plot depicted considerable overlap in morpho-space, indicating a high degree of morphological congruence. This overlap challenges the current subspecies classification based on size and shape traits. Contrary to the proposed latitudinal cline in size, our data indicate that the specimens from the western South America (WSAm), encompassing the Pacific regions of Colombia and Ecuador, and NEC/NWV are similar in size to those from Central America. This finding is consistent with zoogeographical studies that have identified a closer affinity between the bat fauna of the western Andes and Central America Koopman (1976); (1978); (1982); Hoffmann and Baker (2003); Clare (2011). Such patterns suggest the influence of historical events, ecological processes, and landscape heterogeneity (Manel et al. 2003) in shaping the distribution and morphology of bat species, including *Trachops*. The role of heterogeneity of habitats and environmental conditions as causes of morphological and genetic divergence (Turmelle et al. 2011; Lindsey and Ammerman 2016) is also evident in our findings.

### On the subspecific status of Trachops cirrhosus coffini

Despite previous assertions, notably by Fonseca (2019), our findings indicate that *T. c. cirrhosus* and *T. c. coffini* lack distinct morphological synapomorphies and do not exhibit reciprocal monophyly, as required for subspecies designation. This observation aligns with the criteria outlined by Patten (2015), who mentions that a subspecies should be morphologically distinct and geographically circumscribed, yet not necessarily forming a distinct genetic clade. According to Molinari (2023), subspecies designation is appropriate for populations with significant and heritable morphological differences, even if genetic differentiation is insufficient for species-level recognition. Our analyses, however, reveal that the morphological distinctions traditionally used to separate *T. c. coffini* and *T. c. cirrhosus* do not meet these standards. The lack of clear geographical circumscription, combined with our findings that these groups are not morphologically diagnosable nor genetically distinct, challenges the validity of their current subspecific status.

The reliance on size-only differences is contentious. Molinari (2023) underscores that diagnostic morphological characters should not be attributed to phenotypic plasticity. Considering studies indicating size variation as a source of morphological plasticity in bats (McLellan 1984; Jarrín-V et al. 2010; Jarrín-V. & Menéndez-Guerrero 2011; López-Aguirre et al. 2015), the use of body size as a sole criterion for taxonomic decisions becomes problematic. Geographic variation in body size, influenced by a multitude of factors, including genetic adaptations and environmental conditions affecting growth rates, further complicates its reliability as a distinguishing feature (McLellan 1984; Berry et al. 1987; Ebenhard 1990; Jarrín-V et al. 2010; Jarrín-V. & Menéndez- Guerrero 2011; López-Aguirre et al. 2015). Additionally, methodological inconsistencies, including instrumental and human errors, can lead to data discrepancies (Fox et al. 2020), casting doubt on the robustness of size-based differentiation. Our study, therefore, suggests that the current classification of *T. c. coffini* as a subspecies of *Trachops cirrhosus* lacks support, both morphologically and genetically.

### On the specific status of T. ehrhardti

*Trachops ehrhardti* should be elevated to the species status based on the presence of clear diagnosable morphological characters, geographically circumscription in the Atlantic forests of South Brazil (Mata Atlántica), and substantial molecular divergence. Broader cranial measurements (BB, MPW, ZB) allow *T. cirrhosus* to be separated from *T. ehrhardti* and both species form reciprocally monophyletic clades separated by a genetic divergence of over 8%. It is its geographic isolation that would primarily imply loss of ecological exchangeability, and possibly mating impediments with *Trachops cirrhosus*. Originally proposed by Haffer (1969), the Refuge Theory for the Neotropics suggests that the Pleistocene glaciation cycles created contraction and subsequent expansion of forested areas that, in turn, would create allopatry between populations of the same forest-dwelling species, leading to intraspecific differentiation and consequently speciation. Paleopalynological research indicates the high dynamism of Neotropical forested regions: the Atlantic Forest and the Amazon were connected in the past (Vivo 1997), separating themselves as increasing aridity in the Tertiary triggered the formation of the belt of xeromorphic formations between them (Martins et al. 2009). There seemed to be a predominantly arboreal vegetation during most of the Pleistocene, with typical Amazonia and Atlantic Forest tree species found in what it is now the dry diagonal that separates these two biomes. The extent to which these climatic fluctuations and associated vegetation changes affected the patterns of distribution and diversification of the fauna remains a central question in understanding the evolution of forest-associated taxa.

### Implications to conservation

Recognized bat diversity has increased due to new species descriptions and taxa raised from subspecific level or synonymy (Burgin et al. 2018), but also as a result of clarification of cryptic species within several genera (e.g. *Platyrrhinus* (Velazco et al. 2023); *Glossophaga* (Calahorra-Oliart et al. 2021) or *Sturnira* (Yánez-Fernández et al. 2023). These changes in taxonomy may have an impact on the conservation of species. The genus *Trachops* is composed of two monotypic species: *Trachops cirrhosus* maintains its “Least Concern” conservation status at the global level, with no major threats known throughout its range (Miller et al. 2015). However, it is important to revise the status of *Trachops ehrhardti*. The current range of *T. ehrhardti* is restricted to southeastern Brazil. The Atlantic Forest is one of the world’s leading biodiversity ’hotspots’; these areas possess the highest concentrations of species endemism, and simultaneously, are the most severely threatened by habitat loss (Baptista & Rudel 2006). In this biome, the persistent land cover is made up of 476 000 Km^2^, which comprises 9% of the Brazil’s land area (Souza et al. 2020), but is highly fragmented by roads and urban centers, and immersed in a large agriculture matrix, resulting in less than 12% of old secondary forest cover (i.e., >30 years) (de Rezende et al. 2015; Souza et al. 2020). If this scenario remains unchanged, then elevating *Trachops ehrhardti* to a species should trigger immediate protection efforts in conservation.

We present a revised description of the genus *Trachops* and the subspecies *Trachops cirrhosus* and *Trachops ehrhardti* below.

## Taxonomy

The genus *Trachops* can be easily recognized by their highly specialized warty outgrowths or protuberances on the lips and chin. The margin of the nose leaf is finely serrated on the edge, anteriorly connected to the upper lip (Spix 1823). Ears are large and anteriorly covered with hairs projecting beyond anterior margins (Goldman 1925); folds inside the pinnae are well marked. The tail is short and appears on the upper side of a broad interfemoral membrane (Goodwin and Greenhall 1961). The ventral fur is characterized by a pale brown coloration (Spix 1823). Dorsal fur is cinnamon-brown, varying to darker shades in some specimens (Goodwin and Greenhall 1961). The base of the hairs is always white, and the tips are ashy. Underparts are dull brownish tinged with grayish brown (Goodwin and Greenhall 1961).

### Trachops cirrhosus Gray, 1847

*Vampyrus cirrhosus* Spix, 1823:64. No type locality is stated in Spix’s description, but on page 53 Spix said the bats were collected in Brazil. Type locality restricted to Para, Brazil, by Husson (1962: 115).

*Ph[yllostoma*]. *cirrhosum*: Fischer, 1829:126. Name combination.

*Vampyris cirrhosum*: Gray, 1847:481. Emendation of *Vampyrus cirrhosus* Spix

*Trachops fuliginosus:* Gray, 1865:14. Type locality "Pernambuco," Brazil (= *Vampyrus cirrhosus* Spix).

*Phyllostoma angusticeps* Gervais, 1856: 47. Type locality “province de Bahia, au Brasil."

*Tylostoma mexicana* Saussure, 1860 :484. Type locality "les régions chaudes du Mexique."

*Trachyops cirrhosus*: Dobson, 1878:481. First use of name combination and incorrect subsequent spelling of *Trachops* Gray, 1847.

*Trachops coffini* Goldman, 1925:23. Type locality "Guyo, Petén, Guatemala." Type locality restricted to "El Gallo, 8 mi. west Yaxhá, on the Remate-El Cayo trail, Petén, Guatemala" by de la Torre (1956:189).

*Trachops cirrhosus coffini*: Felten, 1956a:189. Type locality restricted to Para, Brazil, by Husson (1962: 115).

This species is monotypic.

#### Amended distribution and habitat

This species is widely distributed in eastern and southern Mexico, extending from Veracruz, in the southeast, southeastward through Central America, into South America, including Colombia, Venezuela, the Guyanas, Ecuador, northern and central Peru, Bolivia, Trinidad, as well as northern, southern, and central Brazil (Figure 7). This species occurs at elevations from sea level (e.g., Belem, Para, Brazil) up to 1800 m (Las Tolas, Pichincha, Ecuador; (Arcos et al. 2007). Records of *Trachops cirrhosus* are predominantly associated with habitats in humid tropical forests, as well as sub-montane and montane forests.

**Figure 7.**
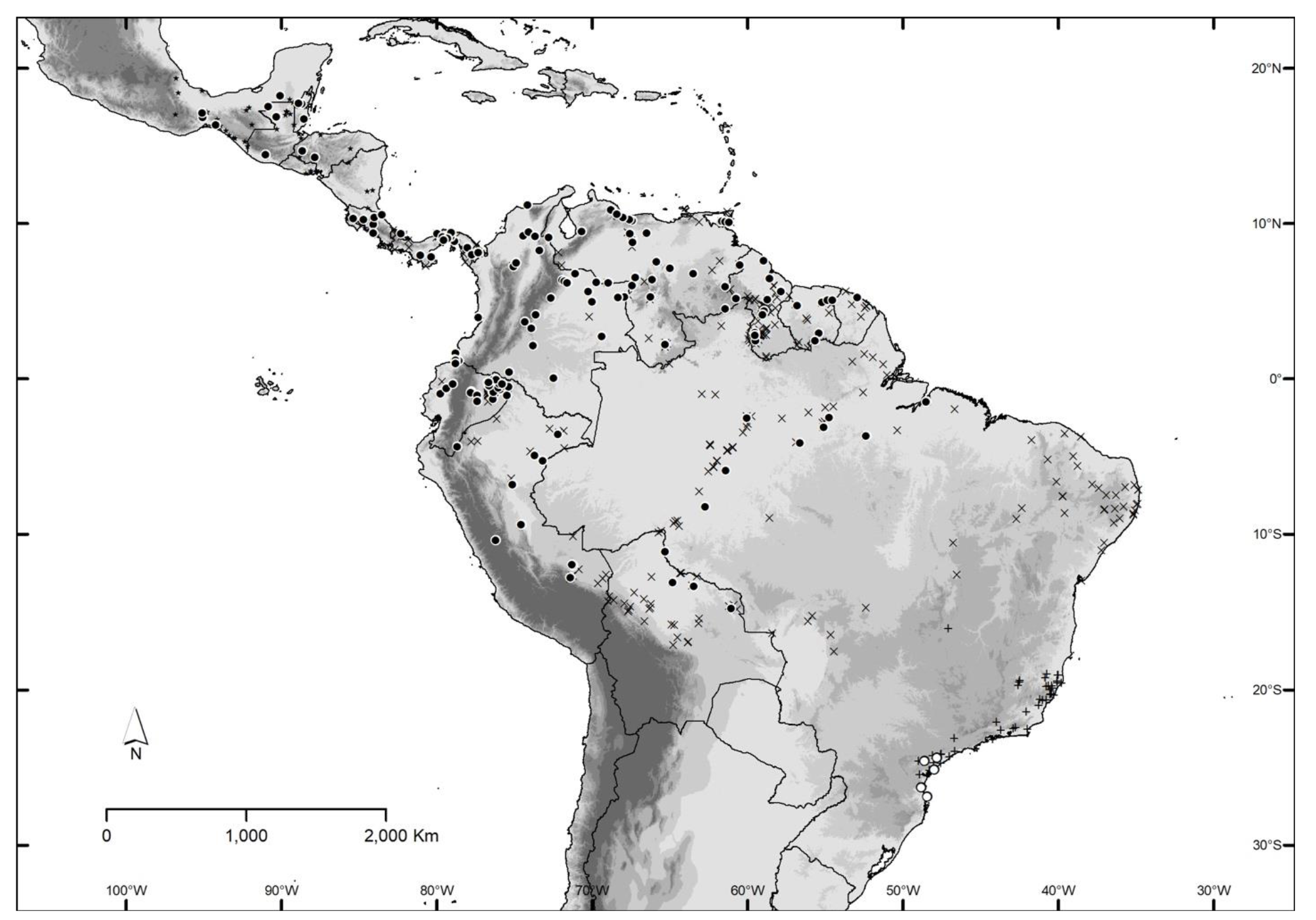
Emended distributions of *Trachops cirrhosus* and *Trachops ehrhardti.* Black circles and crosses (• and ×) represent the recorded locations of *Trachops cirrhosus*. White circles and plus signs (ο and+) correspond to the recorded locations of *Trachops ehrhardti*. The crosses and plus signs are the data collected by Fonseca (2019), sourced from https://github.com/bsf07/Defesa.git.

#### Emended diagnosis and comparison

*Trachops cirrhosus* is a relatively robust (Spix 1823), medium-size bat (FA: 49.57–64.70 mm; GLS 24.70–31.24 mm). Overall size is larger than *Trachops ehrhardti* (Table 5). The skull is large and elongated. The elevated braincase above the rostrum was previously believed to be a distinguishing characteristic, potentially separating the subspecies *cirrhosus* and *coffini*. The latter was described as having a smaller and slenderer shape compared to *T. c. cirrhosus*, although similar to *T. c. ehrhardti*. This characteristic is not diagnostic; rather, it displays variability across the entire species distribution. The sagittal crest may or may not be developed in males and females, contrary to the well-developed sagittal crest character mentioned by Goodwin and Greenhall (1961). Poorly developed sagittal crest is common in most specimens reviewed for the entire area of distribution. However, the faint notch in the cutting edge of the upper incisors (Goldman 1925), is variable. Inferior premolar 1 (p1) is taller compared to the same tooth in *T. c. ehrhardti* which is wider overall. The first lower premolar is three-fourths the height of the third premolar but wider in cross-section (Dobson 1878). The mandible of *T. cirrhosus* is broad and robust; upper and lower premolars are relatively narrower than in *T. ehrhardti*. The dentary is tall and has a contracted premolar toothrow, and an expanded molar toothrow, with the last molar being closer to the fulcrum, a high coronoid process, and an expanded angular process (Figure 4).

**Table 5.**
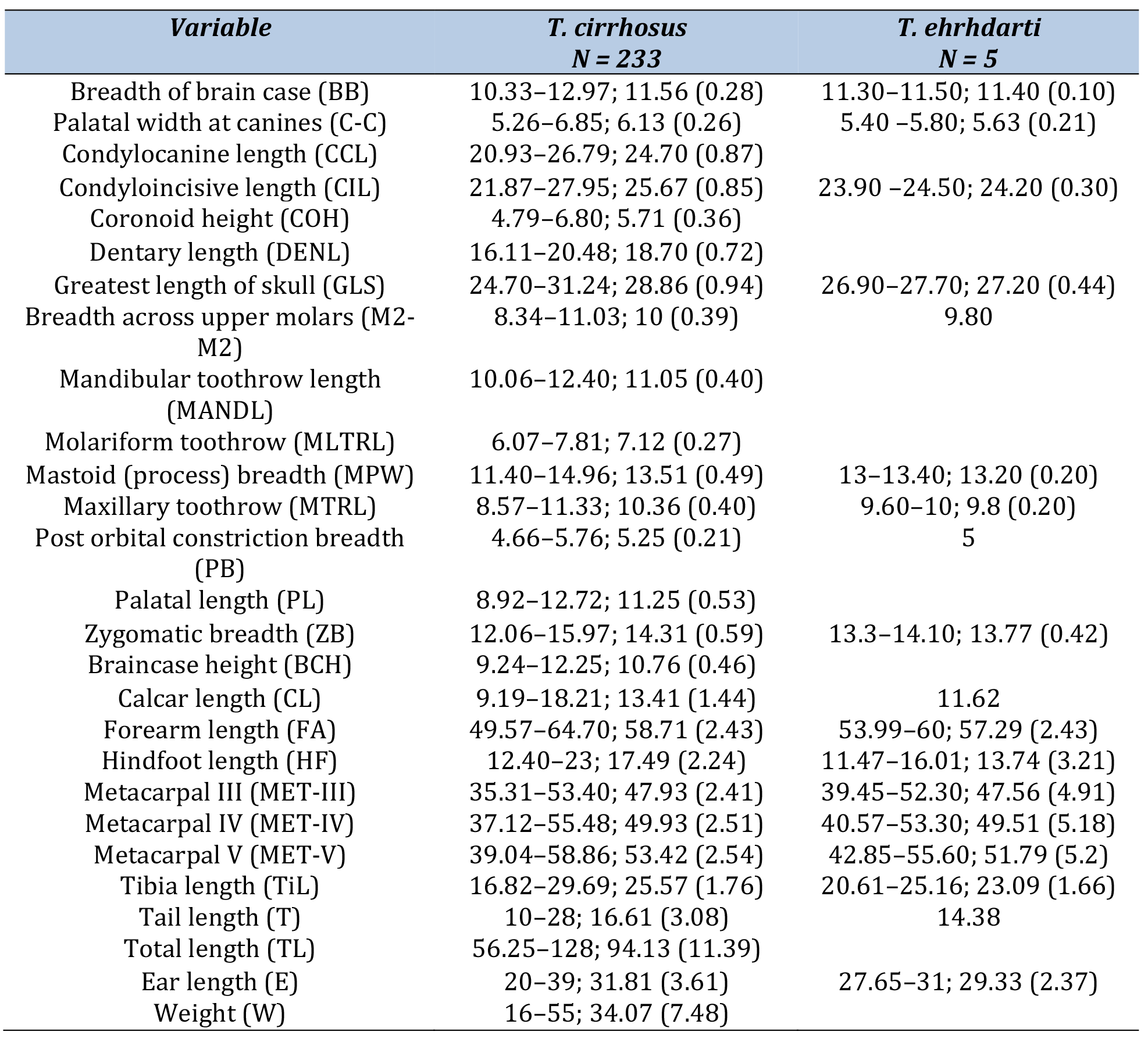
Measurements (mm) and body masses (g) of *Trachops cirrhosus* and *T. ehrhardti*. Descriptive measures: minimum–maximum; arithmetic mean (standard deviation)

#### Remarks

The character of m1 with a less developed paraconid in *T. c. cirrhosus* than in *T. coffini* mentioned by Dobson (1878) was not evident as a diagnostic character.

#### Natural history

*Trachops cirrhosus* is a large-eared gleaning bat that hunts by listening for prey-generated sounds (Obrist et al. 1993). It roosts in caves, hollow trees, road culverts, sewer systems, and buildings, in groups of up to 50 individuals (Hall and Dalquest 1963; Fleming et al.1972; Alvarez-Castañeda and Álvarez 1991; Kalko et al. 1999; Halczok et al. 2018). The relatively sedentary foraging behavior of these gleaners is reflected in their wing morphology, a characteristic that influences their use of small foraging areas and short commuting distances. They have been observed to engage in two distinct flight patterns while foraging: long flights that extend for several uninterrupted minutes and short sally flights lasting less than one minute (Cramer et al. 2001). These bats possess relatively short and broad wings, an adaptation that enhances their maneuverability, particularly in environments filled with obstacles (Marinello & Bernard 2014). However, this wing morphology comes with a trade-off, as it makes continuous flight over extended distances energetically costly (Norberg & Rayner 1987; Kalko et al. 1999).

*Trachops cirrhosus* hunts for prey in continuous flight, presumably depending on prey availability. Prey is usually taken from the substrate (gleaning mode) in a brief landing or may be caught occasionally on the wing (aerial mode) (Kalko et al. 1999). *Trachops cirrhosus* is a carnivorous bat that feeds on a wide variety of prey species, including insects, frogs, lizards, and other small vertebrates (Gardner 1977; Pine and Anderson 1979; Barclay et al. 1981; Tuttle & Ryan 1981; Bonato and Facure 2000; Bonato et al. 2004; Giannini and Kalko 2005; Page and Jones 2016).

Additionally, it has been observed consuming fruits and seeds (Whitaker & Findley 1980; Humphrey et al. 1983;Cramer et al. 2001). Foraging areas of 3 to 12 ha and commuting distances between roost and foraging areas are < 2 km (Kalko et al. 1999). The emergence time and activity peak of *T. cirrhosus* coincided with the maximum calling activity of the Tungara frogs, Leptodactylidae, (Tuttle & Ryan 1981;Ryan et al. 1983). The number and duration of long flights reduce as the calling declines; giving way to an increased frequency of short flights, which might also indicate a switch from frogs to other prey such as insects (Belwood 1990; Kalko et al. 1996, 1999). This bat is very flexible in its responses to prey calls by updating acoustic information with echo acoustic and gustatory cues, as it approaches potential prey, enabling bats to avoid potentially lethal mistakes (Page and Jones 2016). Captive studies show that cues can also be learned socially and transmitted across individuals (Jones et al. 2013; Page and Jones 2016; Flores et al. 2020).

Very little is known about the reproductive biology of *T. c. cirrhosus*. Females give birth to one offspring at a time coinciding with the start of the rainy season (Flores and Page 2017), but the gestation period length is unknown. Females of this species have been reported to be pregnant in April to March and December (Villa-R 1967; Alvarez-Castañeda and Álvarez 1991). In Brazil, Trajano (1984) suggested a polyestrous reproductive pattern, with two annual birth peaks, one before and the other after August. In Trinidad and Tobago, Goodwin and Greenhall (1961) found a colony of *T. cirrhosus* composed of 6 individuals of both sexes, including pregnant females in March (Bredt et al. 1999). The social structure of *T. cirrhosus* is still not fully understood, but a recent six-year study in Panama showed evidence of female philopatry and preferred co-roosting associations in both sexes; kin-biased associations were also detected among pairs of females but not males (Flores et al. 2020).

During the mating season, reproductive males have enlarged testes and create an odorous substance that is smeared on their forearm, called forearm crust. Flores and Page (2017) and Flores et al. (2019) discovered that fringe-lipped males scratch their body with one hind claw, including the area around a prominent mid-ventral chest gland, insert the same hind claw into the mouth, and then repeatedly lick the forearm. Apparently, this substance is not related to female preferences since two-thirds of females selected the scent of a male without forearm crust, but it would play a fundamental role in male-male interactions (Flores et al. 2019).

### Trachops ehrhardti Felten, 1956

*Trachops cirrhosus ehrhardti* Felten, 1956b:369. Type locality "Joinville, Sta. Catarina, Brazil."

#### Distribution and habitat

Southeastern Brazil. Specimens are known from the humid environments of the Atlantic Forest of the states of Santa Catarina, Parana, Sao Paolo, Mina Gerais, Rio de Janeiro and Espiritu Santo (Figure 7).

#### Diagnosis

Medium-size bat (FA: 54–60 mm; GLS 26.90–27.70 mm). Smaller than *Trachops cirrhosus* (Table 5). The ventral fur from the base of the hair shows a lighter shade of brown that gradually transitions to an ashy color at the tips. Dorsal fur is characterized by a reddish-brown to cinnamon-brown hue. The base of the hair appears whitish, creating a contrast with the slightly ashy tone at the tips. Underparts are light brownish with a grayish tint. This coloration is attributed to the presence of light-colored tips on the hairs, and there are white-tipped hairs specifically found on the animal’s underparts. The pinnae are hairy, with marked folds. Lips and chin are ornamented with wart-like protuberances and ears are large and clothed with hairs projecting conspicuously beyond anterior margins as in *Trachops cirrhosus* (Goldman 1925).

Skull is noticeably smaller in comparison to *T. cirrhosus*. Braincase is smaller, forming an angle with a less pronounced rostrum than in *T. cirrhosus*, however, this character shows more degree of variation. Sagittal crest is not developed. In the analyzed skulls, which belong to the type series, it is observed that the lingual edges of the w-shaped stylar shelf are longer. The molar crowns are relatively longer, and in the dorsal view, the parastyle, mesostyle, and metastyle of the second and third upper molars are further away from the maxillary bone. The formed cusps are further apart than in T. *cirrhosus*, giving them a broader appearance. These characters could be important to identify individuals from the eastern areas of Brazil where there could be contact between *T. cirrhosus* and *T. ehrhardti* (Figure 4).

#### Natural history

To the best of our knowledge, precise data on the natural history of this population is currently lacking, although it is expected to share similarities with *T. cirrhosus.* This species inhabits the Atlantic Forest, which encompasses both lowland and montane systems along the Atlantic coast of southeastern Brazil, as well as the contiguous moist-subtropical forest of the Parana basin (Pavan et al. 2016). Similar to *T. cirrhosus*, it is likely that they seek refuge in caves, hollow trees, culverts, and buildings, often in small groups comprising a few tens of individuals. The species might prey on insects, frogs, lizards, and other small vertebrates. Currently, no information is available regarding their reproductive behavior.

#### Remarks

The species was described by Felten (1956b) based on 3 specimens collected in 1908 by W. Ehrhardt at Joinvile in the Brazilian state of Santa Catarina, considerably expanding the southern limit of distribution of the species in South America. The only distinctive characteristic of the subspecies mentioned by Felten (1956b) is its size, notably smaller in relation to animals from the northern part of South America and similar in size to the former subspecies *T. c. coffini*. In 2019, in a degree dissertation from the Federal University of Espírito Santo (see Fonseca, unpublished data), it was proposed, based on an integrative approach to morphological, ecological, and phylogenetic evidence, that *T. cirrhosus ehrhardti* should be elevated to the species category; however, the species has not been formally described nor has the taxonomy been validated.

#### Comparisons

The size and shape of the skull of the two species is similar, but in *Trachops ehrhardti* it is smaller on average. In *T. cirrhosus*, the skull is larger and more elongated than in *T*. *ehrhardti*, and the braincase is more elevated above the rostrum. However, this character shows a degree of variation. *T. cirrhosus* presents a faint notch in the cutting edge of the upper incisors while is broad, with an open groove leading to a distinct notch in the cutting edge in *T. ehrhardti* (Goldman 1925), but this characteristic is also variable. The first lower premolar 1 (p1) in *T. ehrhardti* is wider overall compared to the same teeth in T. *cirrhosus* which is taller than wider. Molariforms toothrow in *T. cirrhosus* is longer than in *T. ehrhardti*, *Trachops cirrhosus* mandible is broader and more robust than in *T. ehrhardti*. In *T. cirrhosus*, the dentary is tall and has a contracted premolar toothrow, and an expanded molar toothrow, with the last molar being closer to the fulcrum, a high coronoid process, and an expanded angular process. This could be associated with a higher bite force (Nogueira et al. 2009). Regarding the width of the cranium, 3 measurements are broader in *cirrhosus* than *ehrhardti* (i.e. Breadth of brain case, Mastoid process breadth, and Zygomatic breadth).

## Acknowledgments

We extend our gratitude to the FSPI–Doctoral Schools Project of the French Embassy in Ecuador, supported by the Ministry of Europe and Foreign Affairs. This work has greatly benefited from the “Investissement d’Avenir” grants overseen by the Agence Nationale de la Recherche (CEBA, ANR-10-LABX-25-01; TULIP, ANR-10-LABX-0041). We are thankful to Marisa Surovy (American Museum of Natural History), Darrin Lunde (National Museum of Natural History), Nicolás Reyes-Amaya (Instituto de Investigación de Recursos Biológicos Alexander von Humboldt), Oscar E. Murillo-García (Universidad del Valle), and Edith Montalvo (Museo de Historia Natural Gustavo Orcés-V.) for providing access to invaluable specimens. We would also like to thank Adam Ferguson and Bruce Paterson from the Field Museum of Natural History, Marie L. Campbell and Joseph Cook from the Museum of Southwestern Biology, Jacqueline Miller and Burton Lim from the Royal Ontario Museum, and Dr. Irina Ruf from the Senckenberg Naturmuseum Frankfurt for their generosity in providing tissue samples. For their work on the mtDNA sequencing and assembly, we express our profound thanks to Alexandra Bialonski and Marike Petersen from the Bernhard Nocht Institute for Tropical Medicine. We also want to acknowledge Ruben D. Jarrín from Pontificia Universidad Católica del Ecuador and Anika Vogel from the Senckenberg Research Institute for their photographic contributions. Santiago F. Burneo deserves a special mention for granting access to specimens under his care at QCAZ and his help with the maps presented on this manuscript. Our sincere appreciation goes to Diego Tirira and the anonymous reviewers whose insights and comments have significantly enhanced this manuscript.

## Appendix 1.

List of specimens used in the present study including voucher identification, GenBank accession number, taxonomic information, and general features of the mitogenome assemblies.

| Museum number | GenBank Accession number | Species | N° of reads | Mean coverage | N° of mtDNA reads | mtDNA length | GC content (%) |
| --- | --- | --- | --- | --- | --- | --- | --- |
| AMNH-15103 | Pending | T. cirrhosus | 4324942 | 104,79 | 29909 | 16584 | 37,58 |
| AMNH-31522 | Pending | T. cirrhosus | 2236998 | 37,01 | 10356 | 16601 | 37,58 |
| AMNH-94538 | Pending | T. cirrhosus | 2151243 | 23,34 | 5747 | 16591 | 37,37 |
| AMNH-129064 | Pending | T. cirrhosus | 2740862 | 79,92 | 22336 | 16595 | 37,58 |
| AMNH-147767 | Pending | T. cirrhosus | 2789364 | 7,54 | 1891 | 16598 | 37,58 |
| AMNH-185889 | Pending | T. cirrhosus | 3336996 | 17,44 | 4212 | 16596 | 37,25 |
| AMNH-185891 | Pending | T. cirrhosus | 4357962 | 101,3 | 22996 | 16625 | 37,3 |
| AMNH-230213 | Pending | T. cirrhosus | 4526938 | 399,14 | 90418 | 16622 | 37,66 |
| AMNH-267935 | Pending | T. cirrhosus | 3494936 | 164,35 | 37044 | 16621 | 37,53 |
| AMNH-272813 | Pending | T. cirrhosus | 5430224 | 147,99 | 33 493 | 16655 | 37,58 |
| AMNH-278476 | Pending | T. cirrhosus | 5304394 | 57,21 | 12932 | 16653 | 37,59 |
| FMNH-72150 | Pending | T. cirrhosus | 7428142 | 7,64 | 1773 | 16578 | 37,35 |
| FMNH-114877 | Pending | T. cirrhosus | 7529998 | 82,99 | 18741 | 16598 | 37,36 |
| FMNH-174887 | Pending | T. cirrhosus | 5776246 | 1 237,67 | 282126 | 16743 | 37,7 |
| FMNH-174888 | Pending | T. cirrhosus | 5550234 | 282,11 | 64164 | 16713 | 37,68 |
| MSB-311655 | Pending | T. cirrhosus | 4271430 | 68,59 | 15555 | 16600 | 37,58 |
| MSB-311656 | Pending | T. cirrhosus | 4543408 | 261,75 | 59681 | 16602 | 37,58 |
| MSB-311734 | Pending | T. cirrhosus | 4944460 | 169,89 | 38553 | 16603 | 37,37 |
| MSB-311736 | Pending | T. cirrhosus | 5215900 | 62,19 | 14095 | 16603 | 37,37 |
| QCAZ-2459 | Pending | T. cirrhosus | 3690346 | 54,62 | 12560 | 16630 | 37,46 |
| QCAZ-4234 | Pending | T. cirrhosus |  |  |  | 16822 | 37,60 |
| QCAZ-9239 | Pending | T. cirrhosus | 3751390 | 157,88 | 35638 | 16623 | 37,47 |
| QCAZ-10617 | Pending | T. cirrhosus | 5240730 | 7,99 | 1829 | 16620 | 37,47 |
| QCAZ-11632 | Pending | T. cirrhosus | 6075978 | 111,02 | 25064 | 16571 | 37,31 |
| QCAZ-13317 | Pending | T. cirrhosus |  |  |  | 16619 | 37,50 |
| QCAZ-13956 | Pending | T. cirrhosus | 5352738 | 29,08 | 6658 | 16621 | 37,35 |
| QCAZ-14029 | Pending | T. cirrhosus | 6361566 | 104,18 | 23674 | 16619 | 37,42 |
| QCAZ-17165 | Pending | T. cirrhosus | 6480538 | 20,02 | 4522 | 16614 | 37,34 |
| QCAZ-18335 | Pending | T. cirrhosus | 4985186 | 83 | 18723 | 16579 | 37,35 |
| ROM-95747 | Pending | T. cirrhosus | 5305082 | 299,46 | 67789 | 16624 | 37,32 |
| ROM-98701 | Pending | T. cirrhosus | 5550346 | 43,24 | 9792 | 16612 | 37,67 |
| ROM-99493 | Pending | T. cirrhosus | 3211086 | 79,6 | 18085 | 16621 | 37,31 |
| ROM-103359 | Pending | T. cirrhosus | 4896612 | 14,83 | 3326 | 16317 | 36,24 |
| ROM-103495 | Pending | T. cirrhosus | 4853726 | 522,78 | 118097 | 16621 | 37,59 |
| ROM-104214 | Pending | T. cirrhosus | 4148188 | 39,36 | 9909 | 16582 | 37,3 |
| ROM-107843 | Pending | T. cirrhosus | 4250814 | 17,87 | 4049 | 16617 | 37,41 |
| **ROM-107908** | Pending | T. cirrhosus | 3634508 | 67,38 | 15210 | 16593 | 37,4 |
| **ROM-107926** | Pending | T. cirrhosus | 3176256 | 169,41 | 38250 | 16602 | 37,41 |
| **ROM-107935** | Pending | T. cirrhosus | 3613554 | 260,21 | 58765 | 16607 | 37,39 |
| **ROM-107945** | Pending | T. cirrhosus | 4380050 | 281,34 | 63704 | 16637 | 37,65 |
| **ROM-115072** | Pending | T. cirrhosus | 6091173 | 10,53 | 2211 | 16577 | 37,29 |
| **ROM-115495** | Pending | T. cirrhosus | 7436128 | 21,57 | 4895 | 16624 | 37,42 |
| **ROM-117523** | Pending | T. cirrhosus |  |  |  | 16749 | 37,6 |
| **ROM-119312** | Pending | T. cirrhosus | 4225778 | 167,44 | 37808 | 16611 | 37,67 |
| **ROM-121044** | Pending | T. cirrhosus | 4673024 | 426,56 | 96388 | 16613 | 37,56 |
| **ROM-121090** | Pending | T. cirrhosus | 4051452 | 46,35 | 10501 | 16622 | 37,65 |
| **ROM-121159** | Pending | T. cirrhosus | 6174610 | 1006,61 | 227410 | 16594 | 37,62 |
| **ROM-121252** | Pending | T. cirrhosus | 4190554 | 197,92 | 44724 | 16615 | 37,69 |
| **ROM-122057** | Pending | T. cirrhosus | 4946730 | 215,59 | 49150 | 16622 | 37,67 |
| **ROM-125132** | Pending | T. cirrhosus | 5841756 | 217,79 | 48522 | 16631 | 37,65 |
| **ROM-125898** | Pending | T. cirrhosus | 5053904 | 26,77 | 6039 | 16589 | 37,51 |
| **ROM-125914** | Pending | T. cirrhosus | 5364036 | 132,02 | 29962 | 16606 | 37,53 |
| **FMNH-94717** | Pending | T. erhardti | 3597194 | 8,72 | 1996 | 16618 | 37,03 |
| **FMNH-141595** | Pending | T. erhardti | 4767214 | 20,62 | 4716 | 16619 | 37,3 |
|  |  | **Average** | 4731900,47 | 138,74 | 36649,90 | 16615,22 | 37,46 |
|  |  | **Standar deviation** | 1242366,47 | 172,72 | 52650,26 | 60,37 | 0,22 |

